# Recurrent introgression and geographical stratification shape *Saccharomyces cerevisiae* in the Neotropics

**DOI:** 10.1101/2024.09.27.615306

**Authors:** J Abraham Avelar-Rivas, Iván Sedeño, Luis Fernando García-Ortega, Jose A Urban Aragon, Eugenio Mancera, Alexander DeLuna, Lucía Morales

## Abstract

**Highlights:**

1. Phylogenetic analyses of *S. cerevisiae* isolates from agave fermentation reveal novel clades within the Neotropical cluster.
2. The Neotropical cluster shows geographical population structure.
3. Multiple pulses of interspecific introgression are part of the evolutionary history of *S. cerevisiae* in the Neotropics

From yeasts to humans, introgressive hybridization significantly influences the evolutionary history of living organisms by introducing new genetic diversity. Strains of *Saccharomyces cerevisiae* worldwide exhibit introgressions from the sister species *S. paradoxus*, despite the average sequence identity between these species being lower than 90%. While *S. cerevisiae* isolates from the Neotropics are known for their high levels of introgression, the evolutionary events leading to the unusually high prevalence of them remain unclear. Here, we sequenced 216 *S. cerevisiae* isolates living in open agave fermentation across Mexico, a habitat at the interface of natural and industrial environments. The genomes of these strains revealed considerable genetic diversity and population structure linked to geographic distribution, which had been overlooked due to undersampling of this megadiverse region. These strains, along with those from French Guiana, Ecuador, and Brazil, form a broader Neotropical phylogenetic cluster that is notably enriched in introgressed DNA. Surprisingly, the origins and conservation patterns of introgressions indicate multiple hybridization events, suggesting an unprecedented scenario of flexible species barriers in this region. Our findings underscore concurrent evolutionary processes—geographical stratification and multiple introgressions—that shape the genomes of a diverse lineage of *S. cerevisiae*. Neotropical yeasts thus provide a natural laboratory for exploring the mechanisms and adaptive significance of introgressive hybridization in eukaryotic genome evolution.

## INTRODUCTION

Beyond its long-established status as the most extensively studied eukaryote in the laboratory, the budding yeast *S. cerevisiae* has emerged as an outstanding model for ecology, population genomics, and evolution ^1–5^. Genomic analyses of thousands of isolates from natural and human-related habitats around the world reveal that the species is structured in clades that broadly correlate with geography and habitat of isolation ^6–8^. These global genomic studies have also shown that most strains harbor genetic material from other *Saccharomyces* species, a phenomenon referred to as genetic introgression. For instance, most *S. cerevisiae* isolates from around the world harbor genes introgressed from its sister species *Saccharomyces paradoxus* ^9,10,6,11,12^, even though these species diverged 4-6 MYA and have an average nucleotide divergence of ∼14% ^13,14^.

The Neotropical region, which extend from northern Mexico to South America, is biogeographically relevant due to its rich biodiversity ^15^. Some of the *S. cerevisiae* strains from this area isolated from anthropogenic and natural environments group in a long-monophyletic branch. This diverged branch, referred to as a complex clade by Pontes et al. (2020), aggregates three groups of strains, namely Mexican Agave (MA), French Guiana (FG), and South American Mix 2 (SAM2). Notoriously, the handful of sequenced genomes available to date indicate that this Neotropical cluster harbors an unusually high and heterogeneous number of introgressions from *S. paradoxus* ^6,7,16,12^. Considering the vast area encompassed by this region, the few available sequenced isolates likely provide an underestimate of genetic diversity, thus limiting our understanding of the species evolutionary history in this part of the world.

The elevated numbers of introgressions observed in *S. cerevisiae* strains from the Neotropical cluster could explain its high divergence when compared to other clades. Although a fraction of the introgressions from the Neotropical cluster has been associated with populations of *S. paradoxus* from the Americas ^6,12,17^, their precise origins remain to be determined. In Mexico, there are reports of *S. paradoxus* in fermentations ^18,19^, including a lineage that coexists with *S. cerevisiae* in the agave distillery environment and that has not been considered by previous studies ^18^. Therefore, this *S. paradoxus* lineage could be a potential donor for introgressions in the tropical Americas. Comprehensive understanding of both the vertical evolution of *S. cerevisiae* and the origins of its introgressions is essential to accurately determine the dynamics of interspecies gene flow in the region.

To explore the evolutionary history and genomic diversity of *S. cerevisiae* in the tropical Americas in the context of their strong signatures of introgression, we sequenced 216 strains primarily isolated from open agave fermentations in Mexico. Previous studies have highlighted the megadiversity in this region ^20^ as well as high diversity of *S. cerevisiae* in agave fermentation ^21–24^, but whole-genome sequencing efforts have only included nine strains, mostly from a single agave fermenting region ^6^. We show that most of the newly sequenced isolates belong to the Neotropical cluster, including new clades where diversity correlates with the geographic origin of the strains. Moreover, by analyzing the patterns and origins of the introgressions in the context of the relatedness of these lineages, we identified multiple introgression events in their evolutionary history. Notably, some of these introgression pulses were unique to strains isolated from agave fermentations. Our refined phylogeny allowed us to illustrate how recurrent episodes of gene flow between two sister *Saccharomyces* species have unfolded in a megadiverse region of the world.

## RESULTS

To provide a comprehensive view of *S. cerevisiae* diversity in one of the most biodiverse regions in the world ^15,20^, we sequenced the genomes of 216 isolates across Mexico (Supplementary Figure S1, Supplementary Table S1). Strains were primarily isolated from traditional agave distilleries. This fermentation environment provides a good starting point to study yeasts diversity, as spontaneous fermentations have taken place for over 3,500 years in a wide variety of ecosystems, latitudes, and cultural contexts ^25–27^. Specifically, 211 of the sequenced strains were isolated from agave fermentations collected through various sampling efforts conducted across the country between the years 1988 and 2021 ^28,21,29,24,30,22,31,18^. Four of the five remaining isolates were from fermentations of other substrates — pulque (fermented raw agave sap), apple cider, cactus pears, and cooked stems of sotol plants (*Dasylirion* spp.)— while the fifth was an uncharacterized commercial yeast strain used in one of the sampled distilleries; all of them were isolated in Mexico (Supplementary Table S1). With the 216 genomes (median depth coverage = 163X), we assessed the phylogenetic position and population structure of the strains in the context of the known diversity of the species and we also analyzed the dynamics and origins of the prevalent *S. paradoxus* introgressions within these populations.

### Agave fermentation strains belong to the Neotropical cluster

To assess the relatedness of the agave *S. cerevisiae* strains with other isolates of the species, we first performed multidimensional scaling analysis (MDS) with 1,264 genomes, including the 216 newly sequenced strains. Most isolates from Mexico clustered next to other strains from the tropical Americas, such as those from the French Guiana (FG), the South American Mix 2 (SAM2), and the Mexican Agave 1 (MA1, formerly named “Mexican Agave”) clades (Figure 1A). The rich diversity of the newly sequenced strains is evident, as strains from the tropical Americas encompass nearly half of the global diversity shown in MDS component 1, depicted by gray points, and nearly the entire diversity in component 2 (Figure 1A). Only eleven strains grouped with other clades, namely the Mixed-origin, Wild North American, or Wine clades. The population-structure (ADMIXTURE, K=3) showed that ∼97% of the agave fermentation strains had a Neotropical ancestral genetic component (Figure 1B), distinguishing strains from the Neotropics (blue) from wild (green) and domesticated (red) isolates from other regions of the world. The ADMIXTURE analysis also indicated that strains from the tequila industry were highly admixed, bearing a combination of the three ancestral components (Figure 1B). All five isolates from non-agave substrates that we sequenced grouped close to the agave-fermentation strains from the Neotropics.

**Figure 1.**
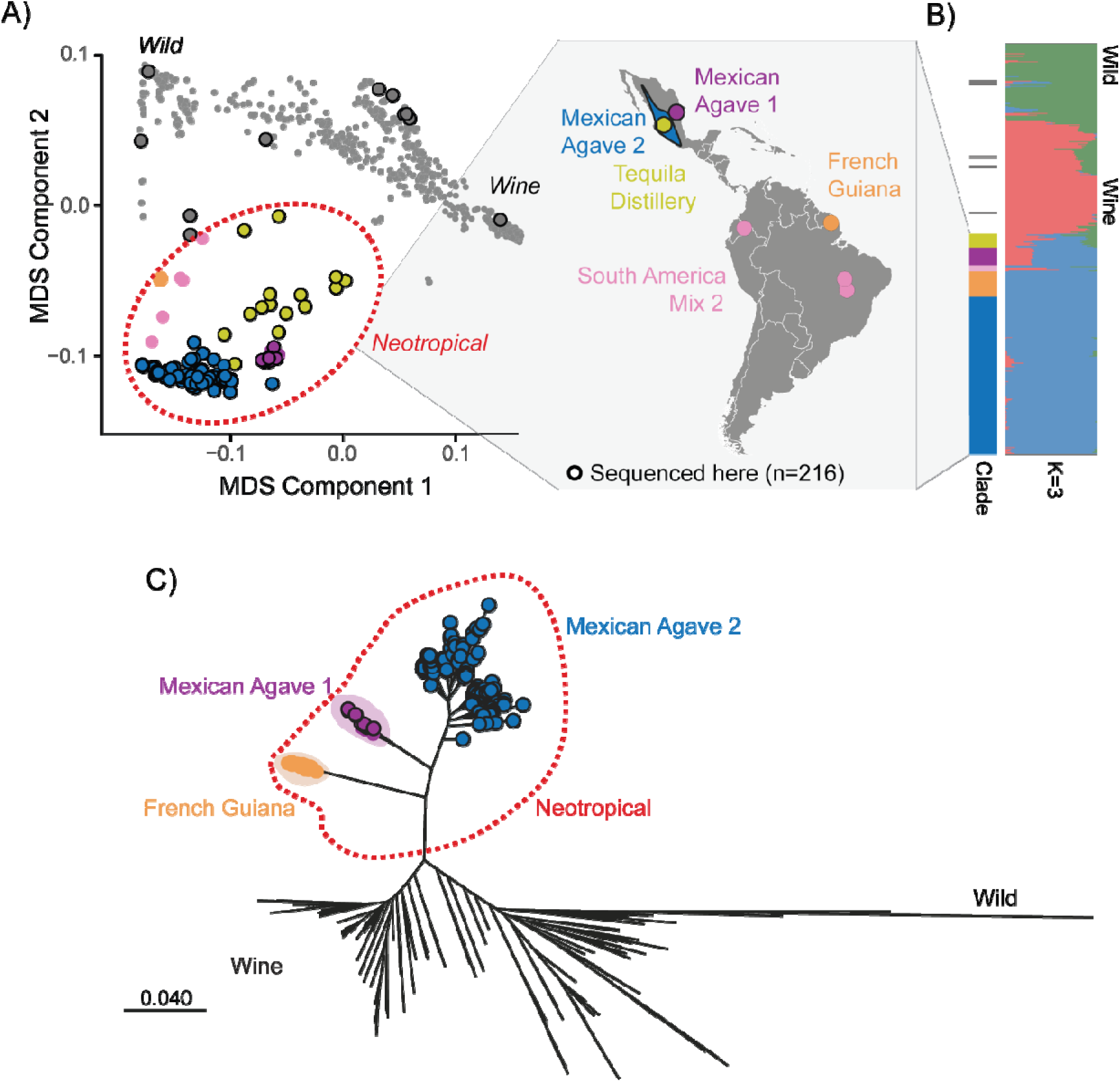
A novel, diverse yeast lineage encompasses most of the diversity in open agave fermentations. **(A)** MDS analysis of 1,264 genomes (173,331 informative sites). Large dots are strains from the Tropical Americas Neotropics, colored according to their clade: “Mexican Agave 1” (purple), “Mexican Agave 2” (blue), “French Guiana” (orange), “Tequila Distillery” (yellow), and “South American Mix 2” (SAM2, pink). Small gray dots indicate isolates from other parts of the world, with the Wine and Wild strains shown as references. Black solid outlines are strains sequenced in this study (n=216). Gray circles with a solid outline indicate the few isolates from agave fermentation that did not group with other Neotropical strains. **(B)** ADMIXTURE population structure at K=3. At the bottom, strains from the Neotropical cluster, with their phylogenetic clades indicated in the first column and map at the left. **(C)** Maximum likelihood phylogenetic tree constructed using 954,449 SNPs in 464 strains including a subset of representative reference genomes and excluding the two admixed clades from the tropical Americas with poor branch support, Tequila distillery, and SAM2.

To establish the phylogenetic relationships of the newly sequenced isolates, we compared them to a reference panel of strains (see Supplementary Table S2). Consistent with the MDS analysis, the phylogeny showed that 205 of the 216 newly sequenced strains (94.9%) were part of a monophyletic group that included the clades MA1, a newly defined clade Mexican Agave 2 (MA2) and FG, plus the two admixed clades Tequila Distillery and SAM2 (Supplementary Figure S2A-B). We refer to this higher hierarchy clade as the Neotropical cluster of *S. cerevisiae*, as nearly all the strains in this group come from the tropical Americas (Figure 1B). Of the remaining sequenced isolates, five strains grouped with the wild North America Oak clade; other five—including the commercial strain— were closely related to the Mixed Origin clade and only one clustered with Wine strains.

The mosaic strains of Tequila Distillery and SAM2 branched with poor support and their presence in the phylogeny changed the relative placement of MA1, MA2 and FG (Supplementary Figure S2). For that reason, these admixed strains were excluded from the main phylogenetic analysis (Figure 1C). In contrast, when SNPS from introgressed segments were removed from the phylogeny, the clades of the Neotropical cluster remained the same. The presence of introgressed regions had a less significant impact on the branch lengths of the diverse Neotropical cluster compared to those of the Alpechin clade (Supplementary Figure S2). Furthermore, the difference in the number of single nucleotide variants was more pronounced for Alpechin than for the Neotropical cluster after the removal of introgression (Supplementary Figure S3).

Three clades could be distinguished among the newly sequenced isolates from agave fermentations in Neotropical *S. cerevisiae*. First, we identified a group of 20 isolates from northeastern Mexico, which included the seven strains previously reported as “Mexican Agave” ^6^ herein referred to as the MA1 clade (Figure 1C). It is worth mentioning that the monophyletic MA1 clade comprised only strains isolated northeast of the Sierra Madre Oriental, a mountain range which runs parallel to the coast of the Gulf of Mexico and acts as a natural barrier, exerting strong influence on the climate and biodiversity on either side of the range ^32,33^. As a result of the comprehensive sampling of the isolates reported here, we identified a novel MA2 clade closely related to MA1 (Figure 1C). Most of the newly sequenced strains in this study (81.9%) belonged to this clade. Two sequenced strains isolated in Mexico ^6^ were previously assigned to the SAM2 clade ^12^ and after these analyses were reclassified within the new MA2 clade. The third clade of agave isolates was composed of 15 strains with heterogenous and admixed genotypes, all sequenced in this study (Figure 1B; Supplementary Figure S2). We refer to this as the “Tequila Distillery” clade, since eleven were isolated from the tequila industry.

Our results show that most strains from open agave fermentations are part of the Neotropical cluster, revealing that what was previously described as “Mexican Agave” represents only a minor fraction of the great genetic diversity found in agave fermentations. Furthermore, the clades within this cluster, including the newly identified groups, rank among the highest in terms of single nucleotide variants when compared to the reference genome, surpassed only by the wild Asian clades (**Supplementary Figure S4**). Together, our phylogenomic and population analyses of *S. cerevisiae* from the tropical Americas bridge the gap of genetic knowledge for the species in this megadiverse region of the world.

### Neotropical yeasts belong to diverse populations stratified by geography

Yeasts from the tropical Americas come from a diverse variety of environments across a broad range of geographical distances. We thus asked to what extent geographic parameters influence the population structure of *S. cerevisiae* in the Neotropical cluster. To this end, we carried out an ADMIXTURE analysis assuming the number of populations that better explained the genotypes according to the cross-validation error (Figure 2A; Supplementary Figure S5). The population structure analysis revealed eleven ancestral components in the Neotropical cluster. Specifically, strains from the MA1, Tequila Distillery, FG, and SAM2 clades each clustered in unique populations, while the remaining seven populations were all from the newly reported MA2 phylogenetic clade.

**Figure 2.**
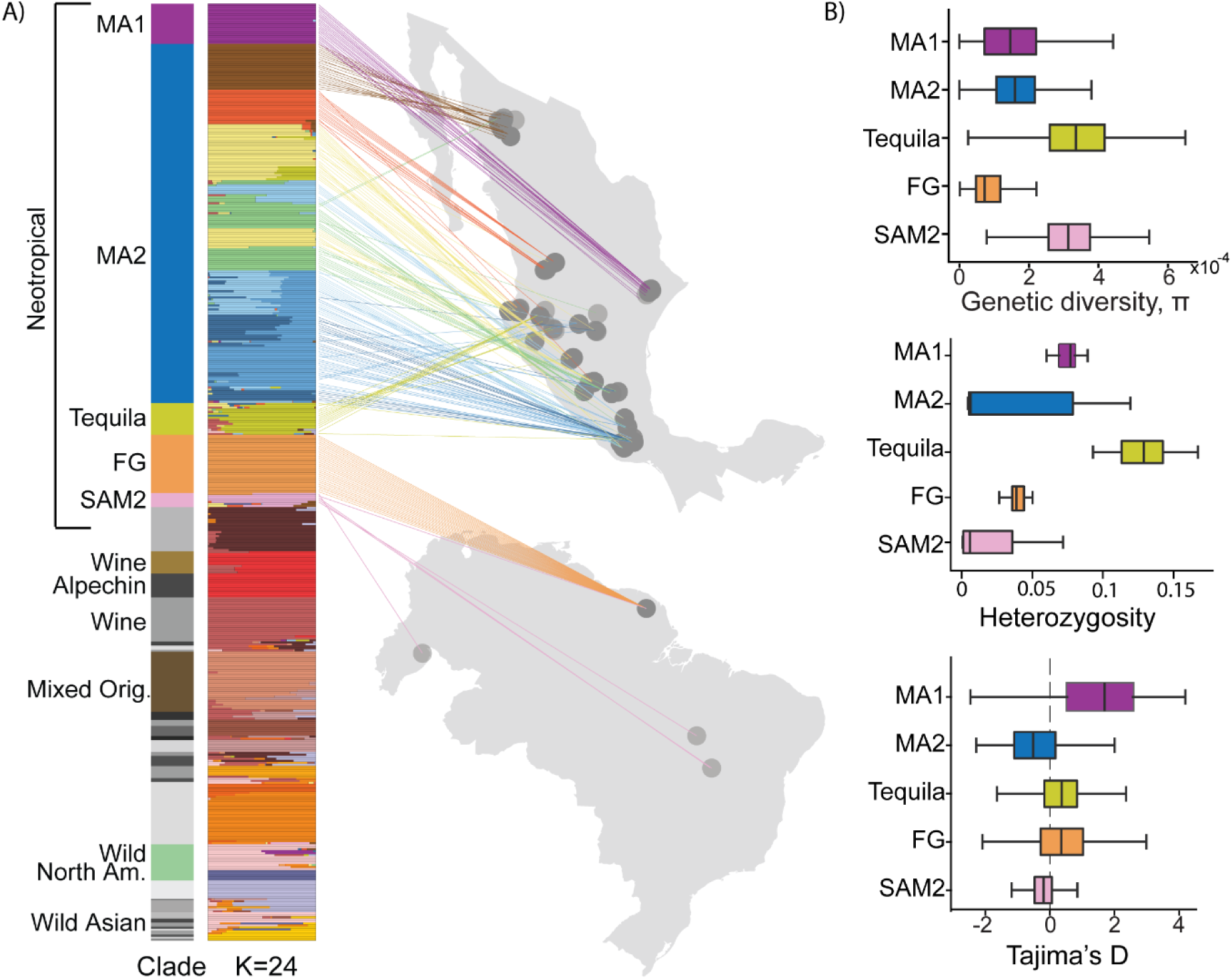
Yeasts from the Neotropics show population structure correlated with geography. **(A)** Population structure of strains from the Neotropics and other regions of the world as reference at K=24 made with ADMIXTURE (n=466); strains are sorted according to phylogenetic clades (first track). To the right, strain’s location is shown in the maps of Mexico (top) and part of South America encompassing Brazil, Ecuador and French Guiana. The colors of the lines connecting strains with geographical location are the same as the most prevalent genetic component of the strain of the Admixture plot. **(B)** Genetic diversity, π (top), heterozygosity (middle) and Tajima’s D (bottom) of clades within the Neotropical cluster.

Given the stratification observed in the Neotropical cluster, we analyzed the geographical distribution of these isolates according to their population structure. The eleven colors in the ADMIXTURE plot representing ancestral components within the Neotropical cluster showed good agreement with their geographic distribution (Figure 2A). This geographic stratification was observed at two levels of resolution. First, each of the five clades of the Neotropical cluster are constrained to specific regions in the map. Second, at the resolution within the MA2 clade, the seven colors in the ADMIXTURE were also stratified by geography. In general, strains from the same agave-distillate producing region grouped together in the phylogeny and had the same ancestral population component. This was also shown by a correlation between geographic and genetic distance in this set of strains (statistic *r*=0.28, *p*<0.05, Pearson Mantel test including MA1 and MA2 strains) and increasing genetic diversity from north to south in the MA2 clade (Supplementary Figure S6). Together, these findings suggest that the genomic diversity in the Neotropical cluster is influenced by geography.

Despite the overall agreement between geography and population structure in the strains from the tropical Americas, there were exceptions suggesting gene flow within and even beyond the Neotropical lineage. For instance, a strain isolated from an open agave fermentation shared close ancestry with wild isolates of the Wild North American clade, while other four isolates closely related to the Wild North American clade showed signs of admixture with different ancestral populations of the Neotropical cluster (Figure 2A). Likewise, there were SAM2 strains showing genetic components of the MA2 and French Guiana clades. Moreover, the Tequila Distillery showed admixture, mostly between different Neotropical populations, but also with other clades such as Wine. Together, these findings suggest that *S. cerevisiae* populations in the tropical Americas have diverged to some extent independently from one another by means of geographic isolation, while other ecological processes or human-related factors have led to gene flow, which also contributes to their genome diversity.

We used the genomic data to further explore the patterns of genetic diversity in the Neotropical strains. The strains from each clade varied significantly from each other in nucleotide diversity (π), Tajima’s D, and heterozygosity (Figure 2B; Supplementary Figure S7), with some of them displaying remarkably high genetic diversity. For instance, the MA1, MA2, and Tequila Distillery clades all show higher genetic diversity than yeasts from the Wine clade (Supplementary Table 2). This held true when comparing with the highly introgressed Alpechin strains, suggesting that differences in genetic diversity cannot be solely accounted for by their introgressions. In contrast, the FG clade showed a homogenous degree of heterozygosity and low genetic diversity. The Tequila clade had the highest heterozygosity and nucleotide diversity, in agreement with trends observed in domesticated beer isolates ^6^, while SAM2 also showed high genetic diversity, but lower heterozygosity, as reported for wild yeast strains ^34^. High diversity in these clades is in part due to their admixed genotypes (Figure 2A), while high heterozygosity specifically in the Tequila Distillery strains may be associated to their adaptation to the semi-industrial context where they were isolated. Taken together, our population-structure and genetic-diversity analyses indicate that yeasts in the tropical Americas constitute highly diverse, structured populations stratified by geography.

### Different distributions of introgressions show recurrent interspecies gene flow

The genomes of *S. cerevisiae* strains within the Neotropical cluster are characterized by an atypically high number of introgressed regions from *S. paradoxus* ^12,17^. To gain insight into gene flow dynamics, we identified the set of introgressed genes (Materials and methods), their location, and length patterns in each clade of the Neotropical cluster (Supplementary Figure S8A). We found that yeasts from MA1 typically exhibited not only more introgressed genes than all other clades, but also the highest variability in this number, ranging from 119 to 364 introgressed genes per strain. Collectively the introgressions in MA1 covered 7.1% of the *S. cerevisiae* reference genome (Figure 3A). We note that the number of introgressed genes in most MA1 strains was within the range observed for the MA2 strains which featured 96 to 160 introgressed genes. Together, all the introgressed genes in the MA2 clade covered 3.2% of the nuclear genome. However, seven strains from the MA1 clade exhibited longer and more heterozygous introgressions than most MA1 strains (Figure 3A; Supplementary Figure S9). Here, we call an introgression as heterozygous when both *S. cerevisiae* and *S. paradoxus* alleles have been retained in a single genome, suggesting a more recent introgression event. The presence of both alleles in several introgressions of the Tequila Distillery clade suggests admixture with a non-introgressed population, which may also explain the lower number of introgressed genes (Figure 3A).

**Figure 3.**
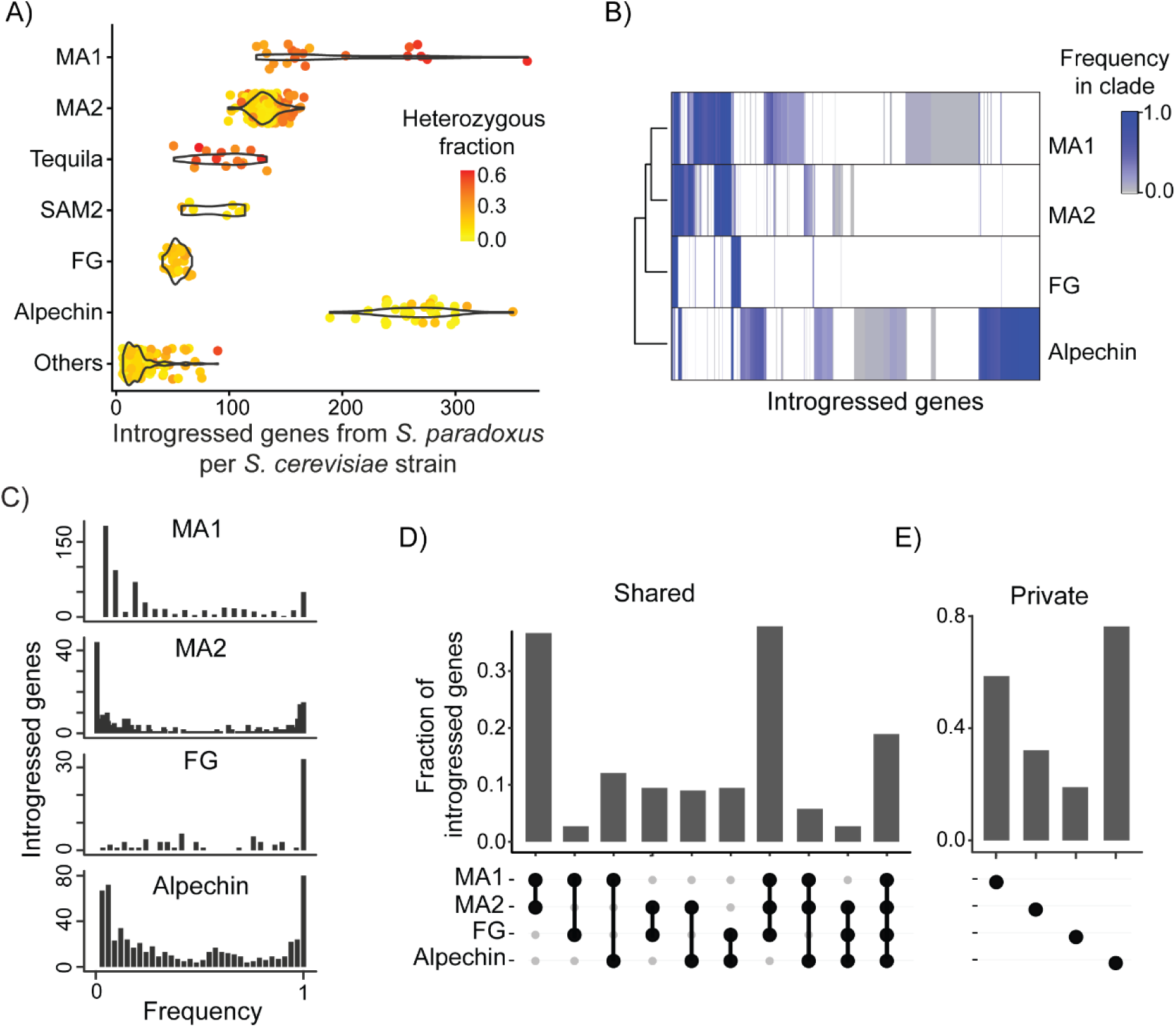
Different patterns of introgressions in the clades from the Neotropics. **A)** Distributions of the number of introgressed genes per strain in each phylogenetic clade. The Alpechin strains are included as a reference of a highly introgressed population. The color of each dot indicates the fraction of introgressed genes that are heterozygous in each strain, meaning that both the *S. cerevisiae* and *S. paradoxus* alleles are present. **B)** Clustering of the introgressed genes found across the Neotropical cluster (n = 1,113). Colors indicate the fraction of strains from each clade that had the introgressed gene. **C)** Distribution of the frequency of introgressions in each clade. **D)** Fraction of the maximum number of possible shared introgressed genes in each comparison. For complete comparisons see Supplementary Figure S10. **E)** “Private” introgressed genes, meaning that they are present only in one of the clades.

All Neotropical clades displayed more introgressed genes than the observed average in other groups from around the world, except for the Alpechin clade. The two clades from South America, SAM2 and FG, displayed fewer and shorter introgressions with a lower degree of heterozygosity, compared to the two Mexican agave clades (Figure 3A; Supplementary Figure S9). This trend suggests that the last introgression pulse of SAM2 and FG likely occurred in the common ancestor of all strains from the Neotropical cluster. These characteristics in the introgression patterns agree with the trends that we observed in terms of the introgressions that are shared, whereby a significant fraction of genes was common between the Mexican Agave clades and, to a lesser extent, with the French Guiana clade (Figure 3B). Importantly, these patterns of shared introgressions were confirmed using an alternative strategy that considered *bona fide* species-specific SNP markers ^12^ for detecting and scoring introgressed regions (Materials and Methods; Supplementary Figure S8).

To further understand how introgressions are shared among clades, we quantified the proportion of strains harboring each introgressed gene within the different clades (Figure 3C). Introgressed genes present at high frequency—at least in 90% of the strains within a/the clade—were common in French Guiana (43.6%) and Alpechin (22.5%). The high proportion of shared introgressed genes within these populations suggests either a strong bottleneck or positive selection acting on introgressed regions. Conversely, introgressed genes present at 10% or lower frequency were more common within the Mexican Agave clades (36% and 45.7%; Figure 3C), which suggests either divergent resolutions of the same hybridization events, recent independent hybridizations in specific strains or a combination of both.

We found that MA1 and MA2 shared the highest fraction of introgressed genes in paired comparisons among clades, suggesting that they were present in their common ancestor (Figure 3D, Supplementary Figure S10). Moreover, the highest fraction in the triple intersection was that of the MA1, MA2 and FG (Figure 3C). These results suggest that a fraction of the introgressions present in the three clades was present in their common ancestor. Additionally, we observed that 14 genes were shared among all four clades, pointing to potential strong selective pressures or artifacts arising from the use of a reference genome lacking those specific genes. In terms of “private” introgressions—those present in a clade but absent in all others—we scored 18.9% in FG, 32.2% in MA2, and 58.6% in MA1 (Figure 3E), while most introgressed genes in the Alpechin reference clade (76.3%) were exclusively found in this clade (Figure 3E). The high percentage of private introgressed genes in Alpechin is due to hybridization events not shared with Neotropical strains ^6,12^. Likewise, MA1 and MA2 isolates exhibited a high fraction of private introgressions, potentially resulting from introgression pulses that are not shared with the related FG clade. Together, the patterns of disparate introgression distribution suggest recurrent interspecies gene flow episodes between *S. cerevisiae* and *S. paradoxus* from the tropical Americas.

### Introgressions in the Neotropical cluster trace to different lineages of *S. paradoxus*

To track the history of the observed interspecies gene-flow episodes, we determined the most likely *S. paradoxus* lineage from which each introgressed gene derived. To this end, we scored sequence similarity of each introgressed block against a panel of *S. paradoxus* genomes including representative strains from the Americas (Figure 4A). Our results suggest that most introgressed genes in the Mexican Agave clades derived from a lineage of *S. paradoxus* that has only been isolated in Mexico and that cohabits in agave fermentations, *SpB_Mx* ^31^. In contrast, the majority of the introgressed genes in the FG strains tracked to *SpB,* a different lineage of *S. paradoxus,* which is not commonly found in agave distillery contexts ^31^. As expected, the introgressed genes in the Alpechin group most closely resemble the European *SpA* lineage, previously identified as the parental population of these introgressions ^35,36,12^. These observations suggest that at least two lineages of *S. paradoxus* have hybridized with ancestors of the extant Neotropical strains, underscoring the close relationship between these two species in the evolution of *S. cerevisiae* in the region.

**Figure 4.**
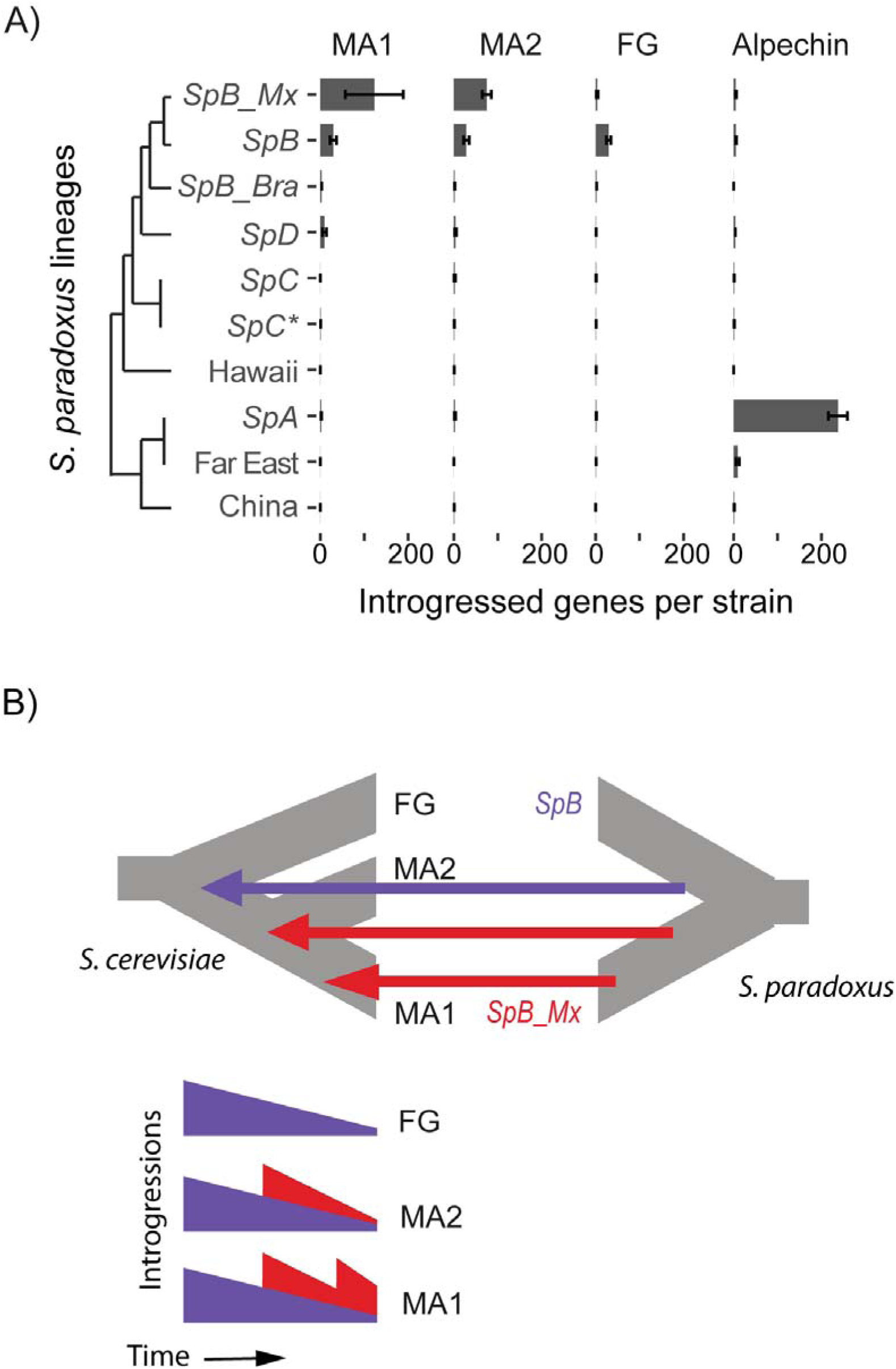
Multiple introgression pulses from two *S. paradoxus* lineages in the Neotropical *S. cerevisiae* cluster. (**A)** The average number of introgressed genes from each lineage of *S. paradoxus* in each clade of Neotropical *S. cerevisiae*. Error bars are the standard deviation. The Alpechin clade is shown as a reference. (**B)** Scheme of the model showing multiple introgression pulses as the most plausible evolutionary scenario for the observed patterns of number and origins of introgressed genes along with the levels of heterozygosity and length of the introgressed blocks.

## DISCUSSION

Comprehensive sequencing of *S. cerevisiae* genomes in megadiverse regions, harboring a substantial portion of Earth’s biodiversity, is essential to understand the evolutionary history and genomic diversity of this prominent model organism. In this study, we sequenced 216 strains from open agave fermentations, providing an extensive population genomics dataset for the Neotropics. Our phylogenetic and population structure analyses revealed multiple interspecies gene-flow events resulting in introgressions within this geographically structured complex clade.

Introgressions impact evolutionary trajectories across the tree of life, from bacteria to humans ^37,38^. These events are also central to the evolutionary history of *S. cerevisiae*, as evidenced by their presence in isolates across the globe ^6,10–12^. In the Americas, several *S. cerevisiae* lineages feature high number of introgressions from *S. paradoxus*, both in wild and anthropogenic environments ^17,7,39,12^, presumably as a consequence of the co-existence of the two species in the region. Here, we provide a broad picture of the evolutionary dynamics of introgression happening in the tropical Americas. In the most plausible scenario, an early introgression pulse occurred in the most recent common ancestor of the Neotropical cluster before its diversification, where genes of the American *SpB* lineage of *S. paradoxus* introgressed into *S. cerevisiae* (Figure 4B). This first pulse was presumably followed by a subsequent introgression event from another lineage, *SpB_Mx*, in the common ancestor of MA1 and MA2. At least one additional, more recent pulse of introgression from the *SpB_Mx* lineage took place in the MA1 clade, which is supported by high numbers of large and heterozygous introgressions in some strains of this clade. It remains to be determined whether these introgression pulses are simply stochastic byproducts of the coexistence of *S. cerevisiae* and *S. paradoxus* in the region, or if they underlie adaptation processes. Wider sampling efforts in other areas, particularly in South America, will likely expand our knowledge of the interspecies gene-flow dynamics in these yeasts.

Our sequencing effort expanded knowledge of the *S. cerevisiae* diversity and biogeography. We uncovered two clades within the Neotropical cluster: MA2 and the admixed Tequila Distillery. Together with MA1, FG, and SAM2, these five clades constitute the Neotropical cluster. Phylogenetic analysis indicated that the MA1— previously designated as Mexican Agave—and MA2 are sister clades, which in turn are the closest relatives of the FG strains, in agreement with the previously reported phylogeny ^40^. The high variability observed in the remaining SAM2 strains indicates that there probably exist other undersampled lineages in the tropical Americas, a diverse and biogeographically complex region (Morrone, 2014). Additional sampling and sequencing in this region would further elucidate key evolutionary events that have shaped the natural history of yeasts, as discussed herein.

Importantly, we found that the novel genetic diversity in the Neotropical cluster is structured and associated with the geographic origins of the yeast isolates. Geographical influence that shaped the genetic diversity was evident across several dimensions: First, a clear correlation was observed between each of the eleven ancestral components of the strains from the Neotropics and their region of isolation. Second, strains from the MA2 clade exhibit increased nucleotide diversity, in a north-to-south gradient. Finally, the genetic divergence between the MA1 and MA2 clades is associated with the presence of the Sierra Madre Oriental mountain range, which is found at the edge where the Neotropical and Nearctic biogeographic regions coincide. This range is known to act as a geographical barrier impacting the distributions of multiple species ^15,32,33^.

Beyond biogeographical factors, human activities may have played a role in shaping these yeast populations, as evidenced by admixture events within and beyond the Neotropical cluster. It remains to be addressed to what extent the observed yeast population structure is driven by natural factors such as the ecology and biogeography of plant reservoirs and insect vectors or by the human dynamics and practices. For instance, the specific local production practices or the remarkable diversity of fermentation substrates, with over 50 species of *Agave* species used for producing spirits ^41^, could be driving population differentiation. With the rising global demand for agave spirits there is a growing risk that producers may abandon traditional fermentation practices. This shift could lead to the loss of the microbial diversity herein described. This work provides a critical framework for developing conservation and management strategies for these fungi, highlighting their value beyond basic research. The agave fermentation system is not only a reservoir of remarkable yeast genetic diversity, but also a unique natural laboratory to gain insights into the interplay of genetic, ecological and evolutionary factors shaping population differentiation, and interspecies introgression.

## MATERIALS AND METHODS

### Genome sequencing and variant calling

DNA was purified with the MasterPure DNA purification kit as recommended by the manufacturer, and it was sequenced using DNBSeq (2×150 bp; BGI, China). Raw reads were quality assessed with fastp V0.20.0 ^42^. For each isolate, the filtered reads were aligned with BWA-MEM v0.7.4 ^43^ to a concatenated species reference genome that includes *S. cerevisiae* S288C, *S. paradoxus* YPS138, *S. mikatae* IFO1815, *S. kudriavzevii* Cr85, *S. jurei* M1, *S. arboricola* H6, *S. uvarum* CBS7001, *S. eubayanus* FM1318, *K. marxianus* DMKU31042, and *P. kudriavzevii* CBS573. Duplicated reads were marked with Piccard 2.6.0 ^44^, and local realignment around indels and variant calling were performed with GATK v4.1.1.0 ^45^. Allele balance information was incorporated in the vcf files with GATK Variant annotator. The called genotypes were filtered to keep only biallelic SNPs with a minimum depth of 5, a QUAL score of at least 30 and SNPs presents in at least 90% of the isolates. Additionally, SNPs were also obtained by mapping the filtered reads only to the *S. cerevisiae* genome to obtain variants that include introgressions (Supplementary Figure S3, Supplementary Figure S4).

### Multidimension scaling analysis

To analyze population structure, the generated genome-wide SNPs data were used for multidimension scaling (MDS) analysis with plink v1.9 ^46^. The data set consisted of 1,264 isolates, including 1,006 from Peter et al. (2018) and 258 from the Americas including the 216 sequenced by us. Only nuclear genome SNPs were considered, and low-frequency or rare variants were excluded to this analysis using a threshold of missing call rate > 1 %. After the application of this filter, isolates with missing genotype > 10% were discarded. The quantitative indices (components) of the genetic variation for each isolate were calculated on the genome wide average proportion of alleles shared between any pair of individuals within the sample.

### Phylogenetic analysis

Maximum likelihood trees were built with genomes sequenced here and references from all over the world with emphasis in American clades. Haplotype sequences were obtained with vcf2phylip v2.3 ^47^ from the nuclear genome variants derived from the concatenated species genome that mapped to the *S. cerevisiae* sub genome (or directly to *S. cerevisiae* reference to include variation from introgressions for Supplementary Figure 2). The phylogeny trees were inferred by raxmlHPC-PTHREADS-AVX2 ^48^ with the GTR+GAMMA parameter and 100 bootstrap replicates. The visualization of the tree was performed using Microreact ^49^. The first two phylogenetic trees were built for 487 isolates including the 216 sequenced here (Supplementary Figure S2). As admixed Tequila Distillery and SAM2 strains changed the topology within the Neotropical cluster, we removed them from the phylogeny presented in Figure 1 with 464 genomes (Supplementary Table S2).

## ADMIXTURE

To run ADMIXTURE 1.3.0 ^50^ the VCF with 466 genomes was curated using Plink 1.9 ^51^ to remove linkage disequilibrium with parameters 50 window size sliding every 5 nucleotides and correlation of 0.5 as previously run in *S. cerevisiae* population genetics ^52^ which led to 618,253 SNPs. Ten seeds were used to run Admixture. The Cross Validation (CV) error was obtained for models with k=2 to K=35 populations. The lowest median CV error K was 24 and therefore the iteration with K=24 and the lowest CV error was used for visualization. Visualization was made in Pong ^53^.

### Estimation of genetic diversity

To calculate genetic diversity (π) and Tajima’s D, we used vcftools 0.1.14 ^54^ and the VCF with variants that mapped to *S. cerevisiae* after mapping to the concatenated reference and filtering high quality biallelic SNPs with less than 10% of missing data. To estimate the per variant heterozygosity, bcftools 1.9 ^55^ was used on individual strain VCFs to calculate the number of heterozygous variants divided by the total number of variants.

### Analysis of Introgressions

To identify introgressed genes, we compared the *S. paradoxus* section of the alignment to the concatenated reference. A gene was considered introgressed if over half of it had a coverage depth exceeding 25% of the median depth of the entire reference. Additionally, it had to be the ortholog with the highest depth or, if not, its depth had to be at least 25% of its ortholog in the *S. cerevisiae* sub genome. We also required a minimum 5% nucleotide-level difference between the orthologs of *S. cerevisiae* and *S. paradoxus*. An introgressed gene was considered heterozygous if the ratio of the depth coverage between the two sub genomes in the competitive mapping was between 0.25 and 4. Nevertheless, we alternatively used a second strategy with the same VCF file to get the positions of a set of *bonna fide* introgression markers and retained introgression blocks of at least five consecutive markers ^12^ to contrast with our mapping-based definition (Supplementary Figure S8). Visualization was made using ChromoMap ^56^.

Once the introgressed genes were determined, consecutive introgressed genes according to the annotation ^57^ were considered part of a block. To determine the origin of the blocks, we aligned the *S. paradoxus* from Lopez-Gallegos, 2023 to the concatenated multi species reference described above and kept only the regions annotated as introgressions in any of the *S. cerevisiae* strains from this work. We computed the paired distances between all introgressed blocks and the *S. paradoxus* genomes using bcftools 1.9 ^55^ and searched for the *S. paradoxus* with the lowest distance to each introgressed block. If a given introgression was most similar to strains from more than one *S. paradoxus* lineage, no origin was assigned to it. All the genes within a given block were considered to be from the same origin.

## DATA ACCESSIBILITY

### Genome sequencing

Genome sequencing data were deposited in the SRA under the project PRJNA1138754. {Reviewer Link: https://dataview.ncbi.nlm.nih.gov/object/PRJNA1138754?reviewer=304775eb7b2b0uud9o0b1aqb6e}

### Custom scripts

Scripts to identify introgressed genes and their origins along with scripts to run the figures of the article are available at: https://github.com/GELab-LIIGH/ScerYeastGenomesMx2024

## ACKNOWLEDGEMENTS

This work was funded by Consejo Nacional de Humanidades, Ciencias y Tecnologías de México (CONAHCYT grants FORDECYT-PRONACES/103000/2020, CB-2016-01/284992, CF-2023-G-695), Fondo de Investigación y Desarrollo Tecnológico del Cinvestav (SEP-CINVESTAV/023), Programa de Apoyo a Proyectos de Investigación e Innovación Tecnológica DGAPA-UNAM (grants IN209021, IN230420 and IN212524), and the UK BBSRC under the Global Challenges Research Fund (GCRF) Growing Research Capability call through the CABANA Innovation Fund (BB/P027849/1). EM was funded by CONAHCYT for a sabbatical stay (I0200/111/2024). LFG-O is a postdoctoral researcher funded by Conahcyt (4133922); IS is a PhD student at UNAM with a graduate scholarship from Conahcyt. We thank Bernard Dujon, Xitlali Aguirre-Dugua and Maria Avila for critical reading of the manuscript. We thank Luis Aguilar (LAVIS, UNAM), Porfirio Gallegos (Cinvestav), Claudio Lopez (Cinvestav), Aaron de Luna (LIIGH), Alejandra Castillo (LIIGH), Carina Uribe (LIIGH), Maritrini Colón-González (LIIGH) and Jair García (LIIGH) for technical assistance. We thank Xitlali Aguirre-Dugua, Luis Delaye, Diego Ortega, Marcela Sandoval and Alicia Mastretta-Yanes for helpful discussions and Manuel Kirchmayr (CIATEJ), Anne Gschaedler (CIATEJ), Maritza Álvarez (CIAD) and Marc Andre Lachance for sharing their strains.

## Supplementary Tables

**Table S1**. The 216 Strains sequenced in this study and associated metadata.

**Table S2**. Strains and metadata used in this study.

---

**Supplementary Figure S1.**
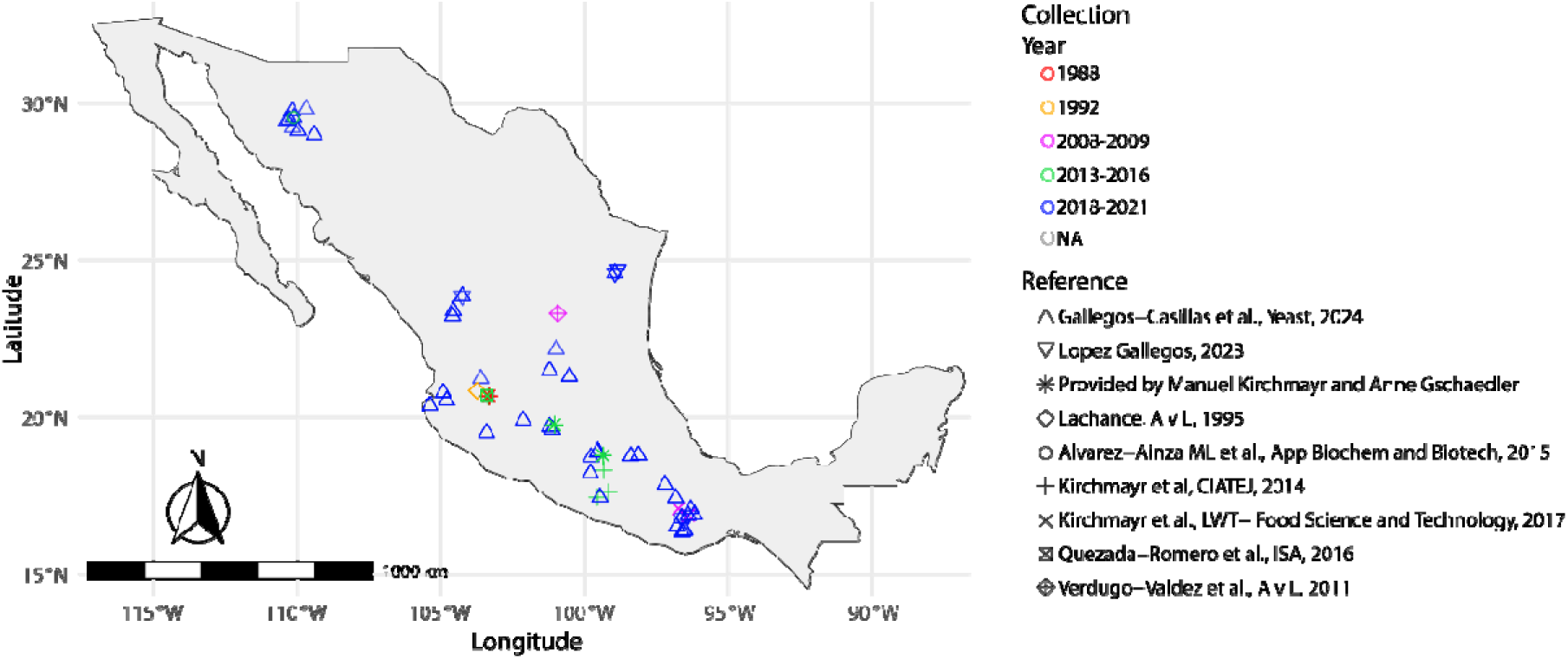
Origin of the 216 sequenced strains.

**Supplementary Figure S2.**
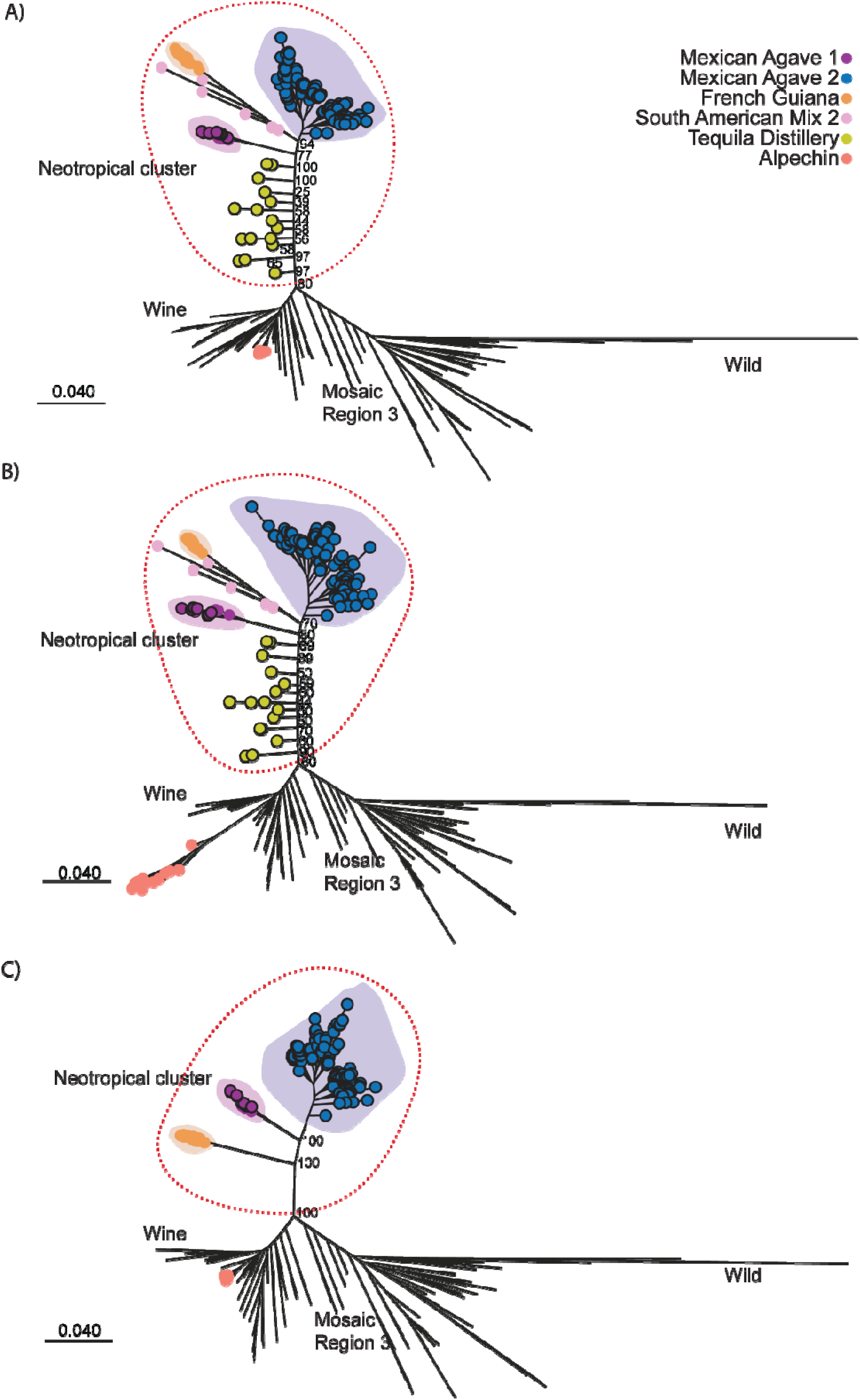
Phylogenies of the Neotropical cluster show the same groups with or without introgressions, but their relative placement is affected by mosaic Tequila Distillery and SAM2 strains. Maximum likelihood phylogenetic trees constructed including a subset of representative reference genomes. (A) Maximum likelihood phylogeny with a complete set of 487 strains and 1,002,285 SNPs excluding introgressions. (B) Maximum likelihood phylogeny of the same set of genomes, including SNPs within introgressions (1,177,709 SNPs). (C) Maximum likelihood of 464 strains as described in the main text. The bootstrap values of relevant branching points show how the Tequila Distillery mosaics hamper reconstruction of the branching order of the major phylogenetic clades. Colored dots at the tip of the branches show the origins of *S. cerevisiae* strains from the Neotropical cluster: Mexican Agave 1 (purple), Mexican Agave 2 (blue), French Guiana (orange), Tequila Distillery (yellow), South American Mix 2 (pink); Alpechin strains are also highlighted (salmon).

**Supplementary Figure S3.**
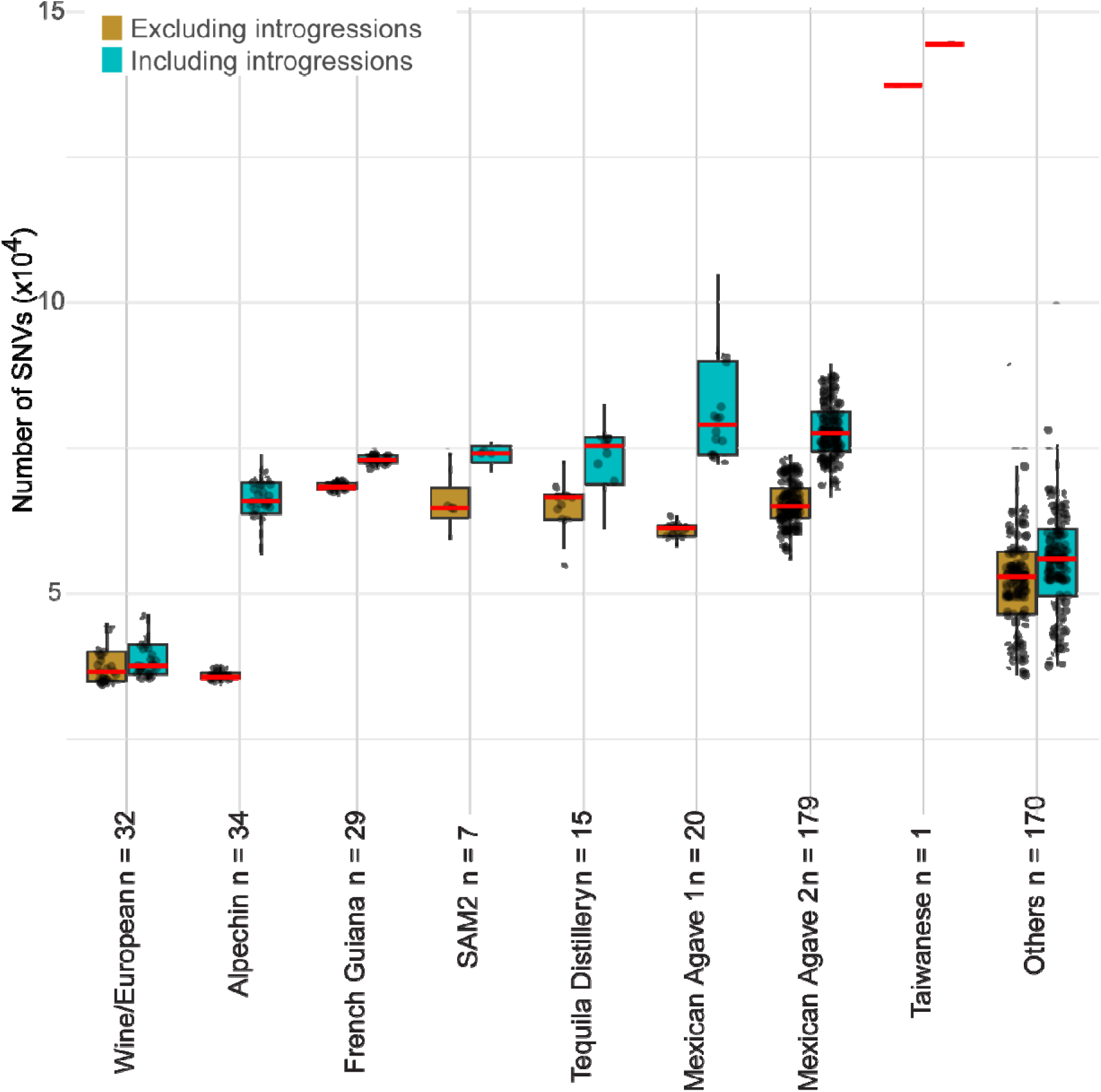
Number of single nucleotide variants of strains with or without introgressions. The numbers are grouped per clade as detected after mapping to the S288c reference genome alone (Including introgressions, blue) or to the multi species concatenated reference as described in Methods (Excluding introgressions, red). Median is shown in red. Wine, Mixed Origin and Taiwanese are shown as reference.

**Supplementary Figure S4.**
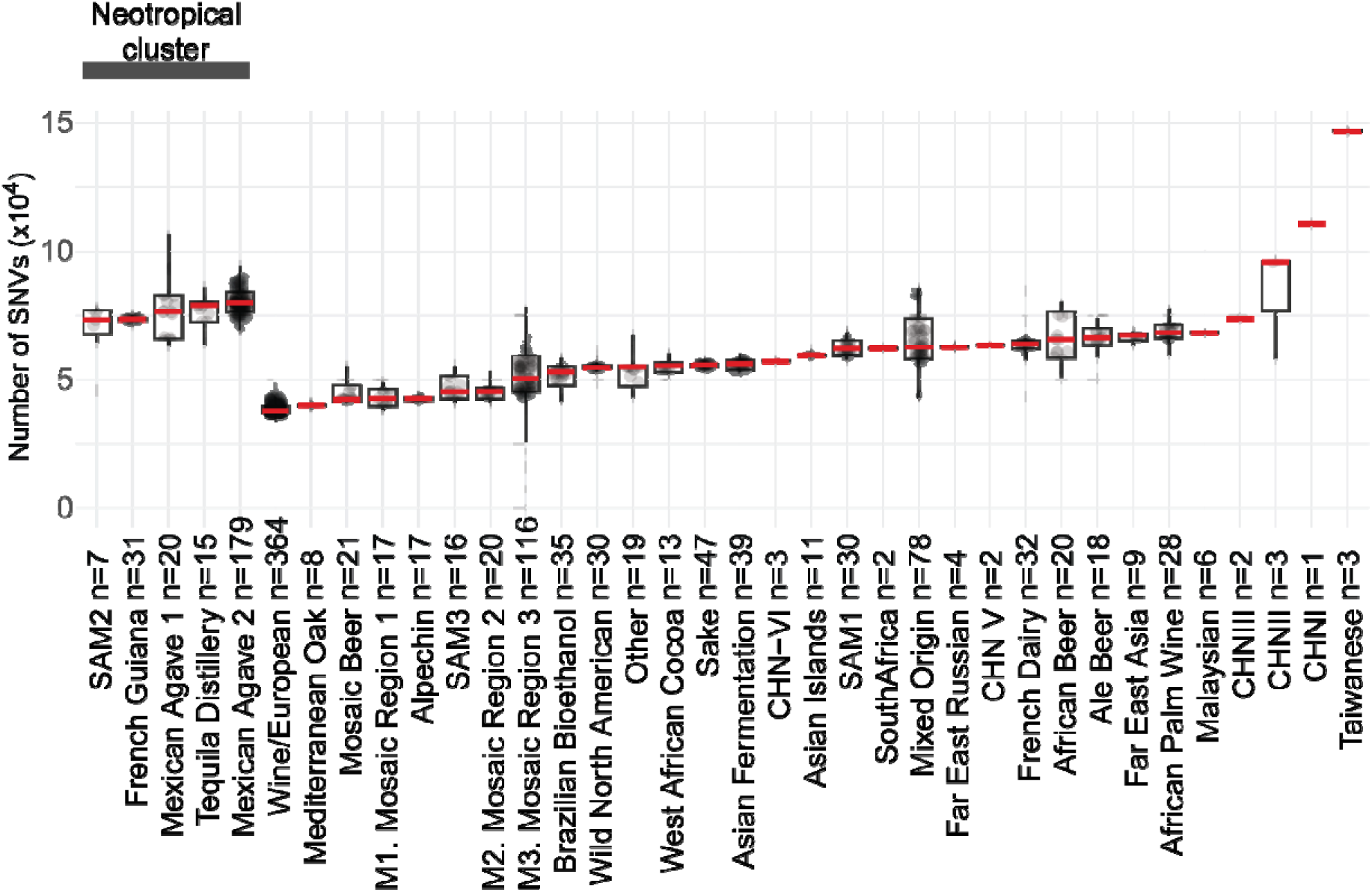
Number of single nucleotide variants of strains representing the world-wide diversity arranged by clade. Neotropical clades are at the beginning and the rest are sorted by the median shown in red. The variants from the newly sequenced strains called as described in Methods were merged with the vcf from Peter *et al.*, 2018, and the number of variants was calculated using bcftools 1.9.

**Supplementary Figure S5.**
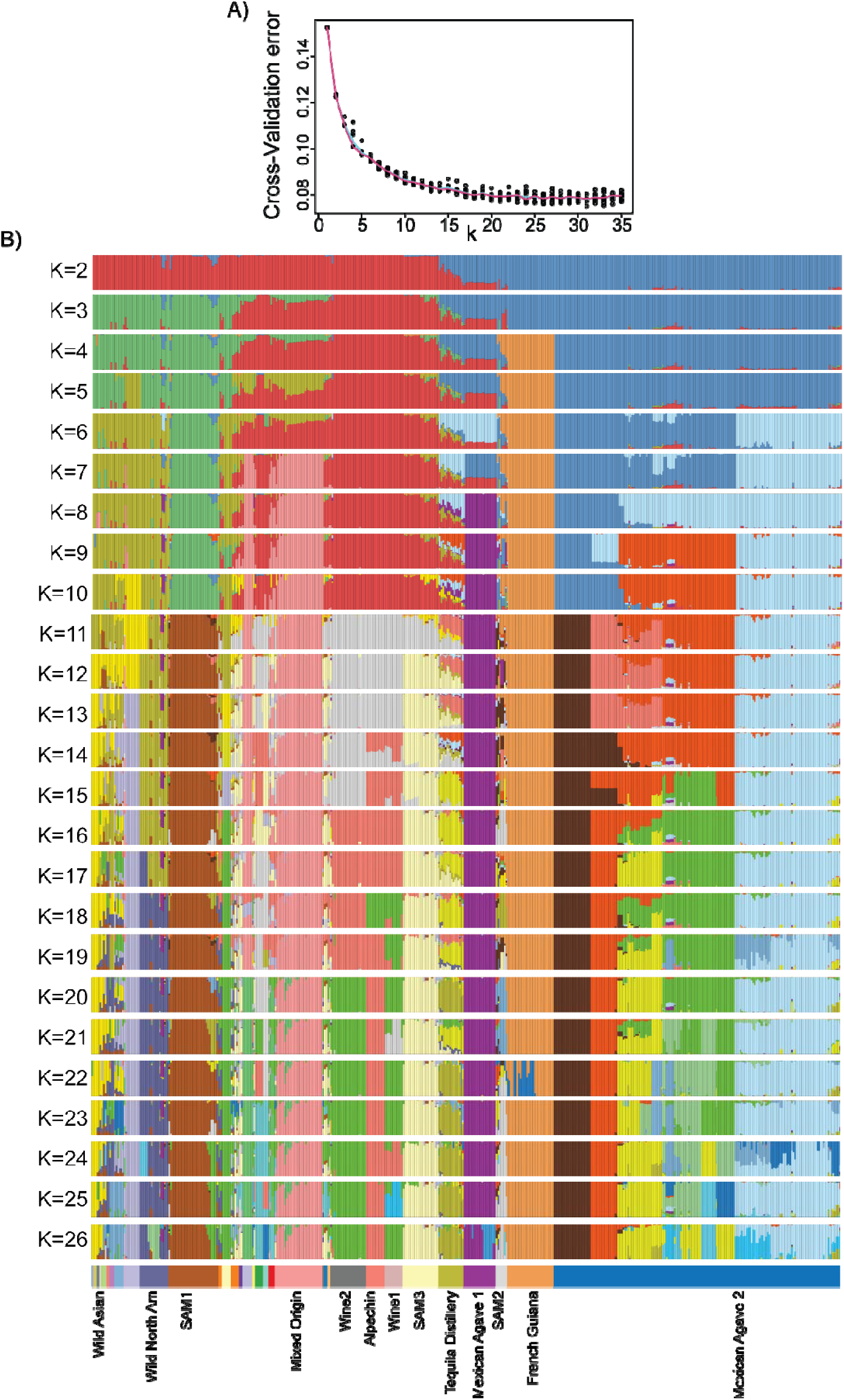
Population structure analysis with ADMIXTURE. (A) Cross-validation error analysis, where each dot is one of ten replicates at different K values. Cyan and magenta lines are the mean and the median, respectively. (B) Admixture population structure analysis of the simulation with the lowest cross-validation errors (K=3 to K=26). South American Mix clades (Tellini 2024) include the following: SAM1 is formed by strains of B1 lineage of Barbosa (2016) and Ecuadorean strains from Peter (2018), SAM2 is formed by B3 lineage of Barbosa (2016) and different unassigned strains from the Americas from Peter (2018), and SAM3 includes lineage B4 of Barbosa (2016), Cachaca strains of Barbosa (2021) and a Brazil spirit strain from Gallone (2016).

**Supplementary Figure S6.**
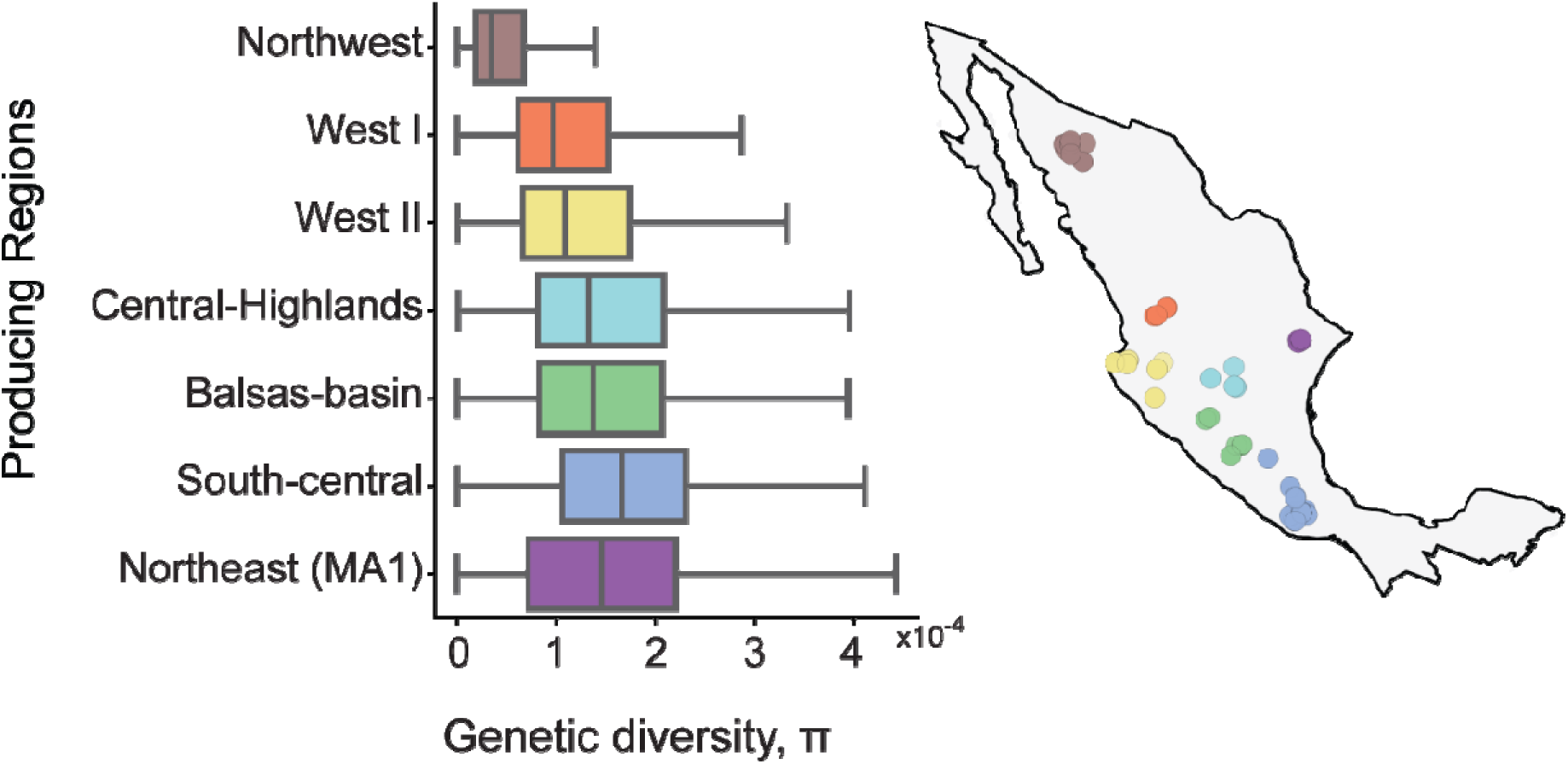
Biocultural regions of agave fermentation for spirit production show a gradient in genetic diversity. Regions are as in Gallegos-Casillas *et al.*, 2024.

**Supplementary Figure S7.**
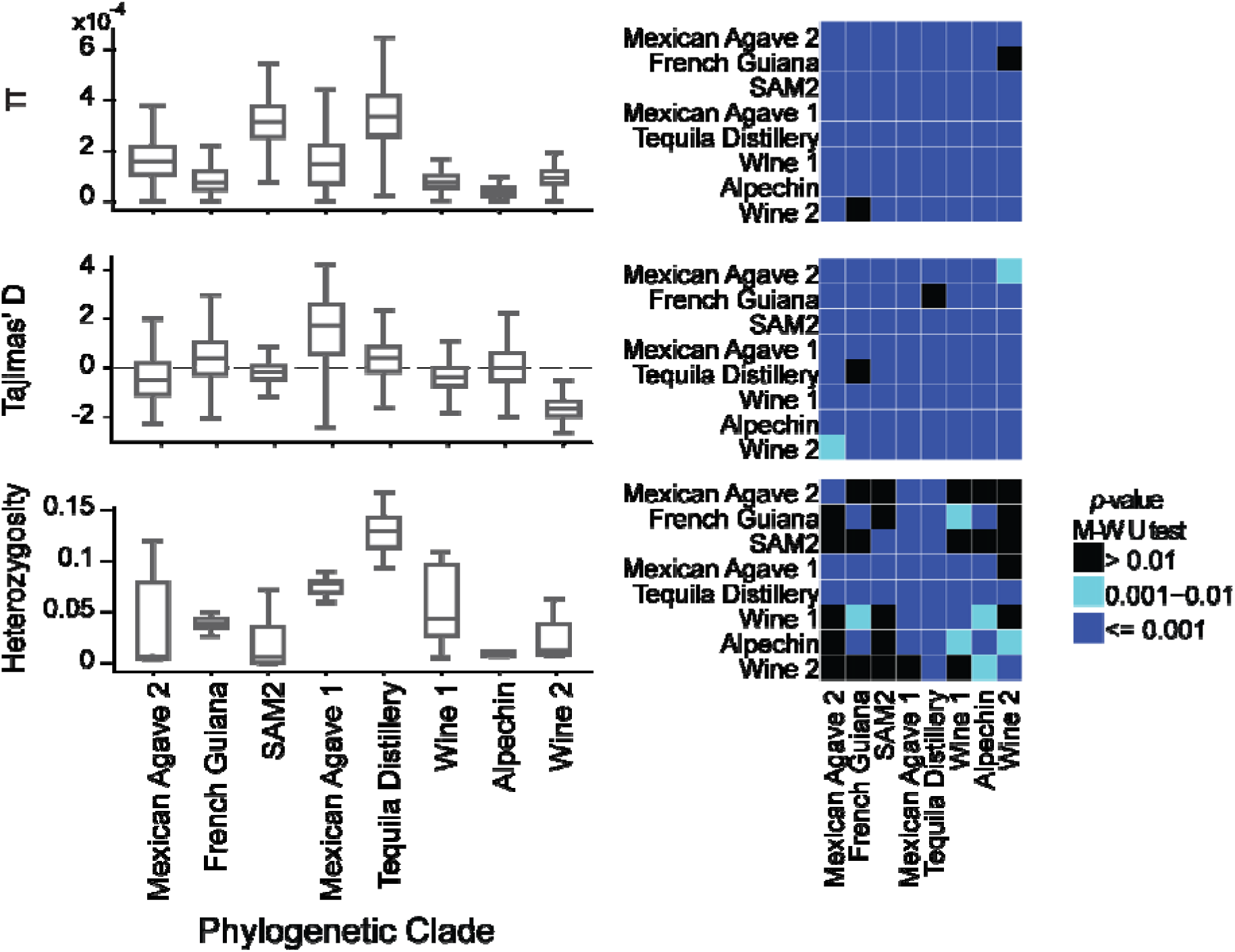
Genetic diversity parameters in clades of yeasts from the Neotropical cluster. Distribution of Heterozygosity (top), Genetic diversity, π (middle), and Tajima’s D (bottom). Each matrix at the right shows the *p*-value of a Mann-Witney paired test.

**Supplementary Figure S8.**
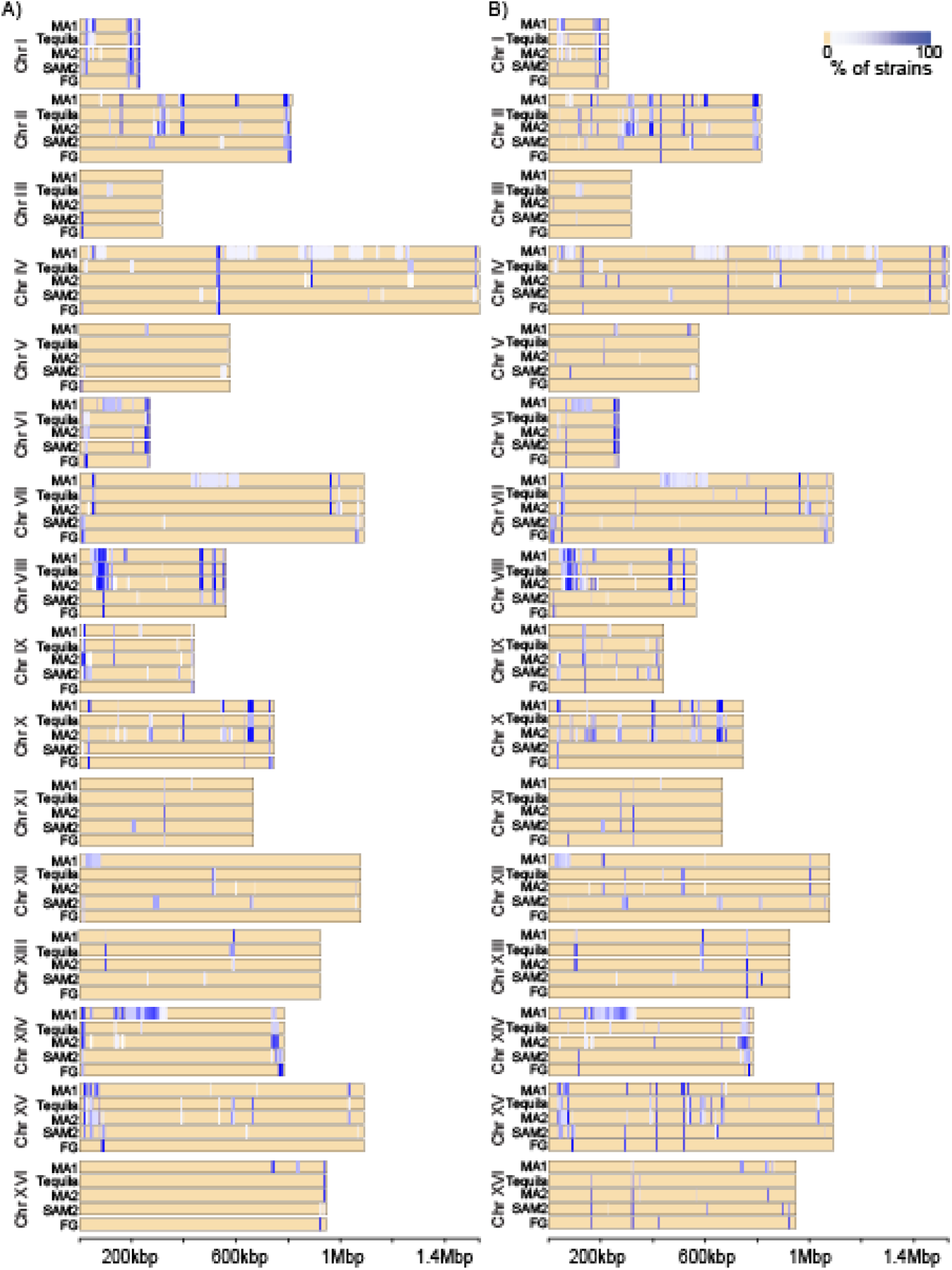
Genomic map of introgressions shows similar distributions regardless of the specific method to define them. For each locus, the color scale indicates the percentage of strains in each phylogenetic clade with introgression, (A) using the coverage of depth after competitive mapping to the concatenated multispecies reference and (B) using the *bona fide* markers from Tellini et al. 2024.

**Supplementary Figure S9.**
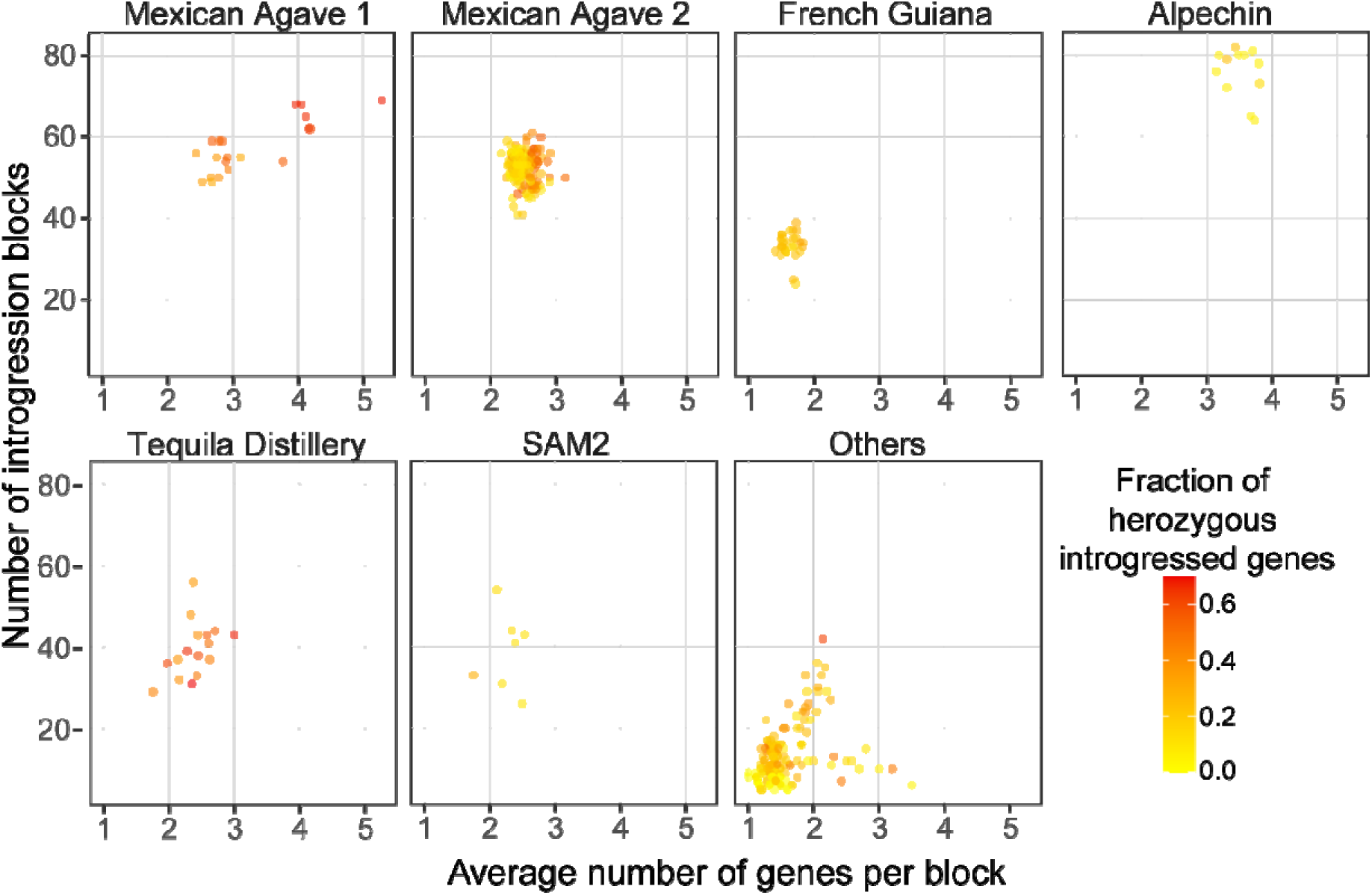
Introgression block size per clade of the Neotropical cluster. Strains with more genes per block tend to also show a higher number of introgression blocks.

**Supplementary Figure S10.**
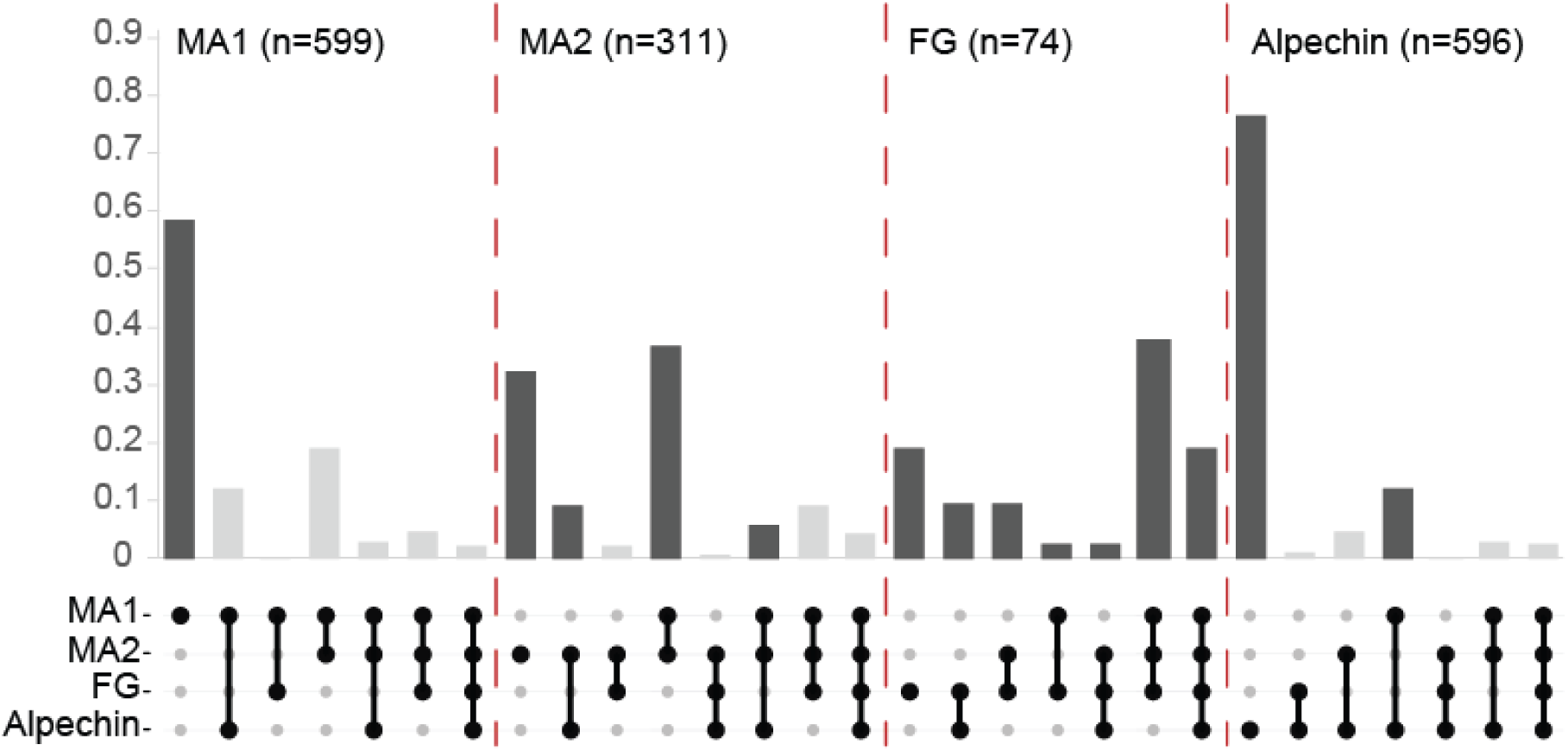
Fraction of shared introgressed genes between highly introgressed clades. The dark gray bars are the same shown in Figures 3D-E. They highlight cases where the fraction is divided by the maximum number of possible shared introgressed genes in each comparison. The maximum number of possible shared introgressed genes is the number of introgressions present in the least introgressed clade.

